# Navigating the translational roadblock: Towards highly specific and effective all-optical interrogations of neural circuits

**DOI:** 10.1101/2020.04.20.049726

**Authors:** Ting Fu, Isabelle Arnoux, Jan Döring, Hirofumi Watari, Ignas Stasevicius, Albrecht Stroh

## Abstract

Two-photon (2-P) all-optical approaches combine *in vivo* 2-P calcium imaging and 2-P optogenetic modulations and have the potential to build a framework for network-based therapies, e.g. for rebalancing maladaptive activity patterns in preclinical models of neurological disorders. Here, our goal was to tailor these approaches for this purpose: Firstly, we combined *in vivo* juxtacellular recordings and GCaMP6f-based 2-P calcium imaging in layer II/III of mouse visual cortex to tune our detection algorithm towards a 100 % specific identification of AP-related calcium transients. False-positive-free detection was achieved at a sensitivity of approximately 73 %. To further increase specificity, secondly, we minimized photostimulation artifacts as a potential source for false-positives by using extended-wavelength-spectrum laser sources for optogenetic stimulation of the excitatory opsin C1V1. We achieved artifact-free all-optical experiments performing photostimulations at 1100 nm or higher and simultaneous calcium imaging at 920 nm in mouse visual cortex *in vivo*. Thirdly, we determined the spectral range for maximizing efficacy of optogenetic control by performing 2-P photostimulations of individual neurons with wavelengths up to 1300 nm. The rate of evoked transients in GCaMP6f/C1V1-co-expressing cortical neurons peaked already at 1100 nm. By refining spike detection and defining 1100 nm as the optimal wavelength for artifact-free and effective stimulations of C1V1 in GCaMP-based all-optical interrogations, we increased the translational value of these approaches, e.g. for the use in preclinical applications of network-based therapies.

**One Sentence Summary:** We maximize translational relevance of 2-P all-optical physiology by increasing specificity, minimizing artifacts and optimizing stimulation efficacy.

## Introduction

*In vivo* 2-P all-optical approaches are based on simultaneous 2-P optogenetic modulations and 2-P calcium imaging *[1]*. They allow for causal interrogations of neuronal networks on single-cell level in a minimal-invasive fashion by using spatially confined light for readout and manipulation. In recent years, 2-P-based methods have been successfully employed for all-optical control of neural microcircuits *in vivo [2-10]*. Using parallel *[2, 7, 9]* or hybrids of parallel and scanning methods *[5, 8, 10-12]*, mainly involving different variants of computer generated holography (CGH), volumetric imaging of hundreds of neurons across cortical layers and simultaneous manipulation of neuronal ensembles with dozens of neurons became feasible *[4, 7, 10, 12]*. Lately, the revolutionizing concept of all-optical physiology has been successfully applied to modulate complex behavior in mammals *[5, 10]* and was extended to closed-loop designs *[8]*. However, although these approaches are advancing rapidly, important aspects concerning specificity and efficacy have not been sufficiently addressed, yet.

Utmost specificity is of crucial importance for applications of all-optical approaches in the context of preclinical research in animal models of neurological disorders. For instance, across different neurological disorders *[13-15]* an early shift of microcircuit activity towards a hyperactive cortical network phenotype has been observed in pre- or asymptomatic phases, and over time these hyperactive network dysregulations itself may lead to neurodegeneration *[15]*. Although these dysregulations are subtle and include complex changes in the activity of single neurons *[16, 17]*, they represent a promising target for preventive, network-based therapy approaches *[18, 19]*. In this case, using all-optical approaches is not farfetched as the combination of 2-P calcium imaging and 2-P optogenetic stimulation would (a.) allow to detect individual hyperactive neurons *[13-15]* and would (b.) allow to modulate their activity in a highly specific manner, e.g. by silencing hyperactive neurons, while leaving intact network components untouched. However, in *in vivo* imaging experiments optical signals are inherently noisy and only indirectly mirroring neuronal activity. Deflections in the fluorescent trace are not only caused by the underlying neurophysiological signal of interest, but can be due to movement or non-physiological artifacts. This is of particular relevance in preclinical studies aiming for development of treatment strategies in animal models. The translation of putatively promising treatment strategies developed in animal models faced tremendous roadblocks and suffered significant setbacks in recent years *[20-23]*. This makes a highly specific assessment of the prospects of network-based therapies a bare necessity. Indeed, prior to the preclinical development of any therapeutic intervention, both the detection and the manipulation of network activity need to be highly specific. On the detection, ultimately, 2-P calcium imaging does not provide a direct readout of neuronal activity, unlike electrophysiology *[24]*. It reports – at the level of microcircuits – the fast elevation of intracellular calcium levels upon action potential (AP) firing *[25]*. Consequently, the first task towards a specific detection of the optical correlate of APs represents the specific identification of these stereotypical calcium transients. The second task towards a specific all-optical physiology represents the avoidance of optical non-AP-related artifacts and the crosstalk between imaging and stimulation. While e.g. movement artifacts strongly depend on the quality of preparation, stimulation artifacts are experimenter-independent, as they are caused by cross-talk between imaging and stimulation channels. A further important issue, which has not been comprehensively addressed, yet, concerns the spectral efficacy of optogenetic control. The spectral range for effective optogenetic control has still not been thoroughly defined. Studies in slice preparations suggest an increase in photocurrents with increasing wavelength, however the maximum wavelength was limited to 1080 nm *[10, 26]*. Until now, no actuator has been tested using wavelengths beyond this range, mainly due to technical limitations.

Here, we propose a step-by-step approach tackling these aforementioned limitations: I) we provide data on the sensitivity of 2-P calcium imaging in terms of specific, false-positive-free identification of AP-related calcium transients, II) we determine the spectral window for artifact-free stimulation, and lastly, III) we assess the efficacy of opsin stimulation beyond 1100 nm.

## Results

### Implementing a highly specific optical detection of neural spiking activity in cortical circuits

For the application of optical imaging of cortical microcircuit dynamics, i.e. the optical correlate of APs, in the context of preclinical research in animal models of neurological disorders, high specificity is paramount. We have developed a program suite comprising different analysis tools *[14]*, which take into account the rather stereotypical dynamics of an AP-related calcium transient such as the dynamics of its rise, its duration, and its decay (see Material and Methods, *[14]*). This is of particular importance, as a mere deflection of the calcium traces from the baseline could very well have non-AP related origins, such as motion or light artifacts. To test for the specificity and sensitivity of its built-in peak detection algorithm, here, we conducted simultaneous juxtacellular electrophysiological recordings and 2-P calcium imaging in the visual cortex of the lightly anesthetized mouse. For that, we performed viral-based gene transfer of the genetically encoded calcium indicator GCaMP6f in cortical neurons. In these bimodal recordings, a stable calcium trace with large-scale deflections could be observed (Fig. 1 A, B). This calcium trace was synchronized with the juxtacellular recordings in voltage clamp mode, displaying typical sparse spontaneous AP firing *[27]*. We set the peak detection criteria in a way that no false-positives, i.e. detection of calcium peaks in the absence of temporally locked APs, was identified (Fig. 1 B). It has to be noted, that indeed, simply correlating relative fluorescent changes would have led to the false identification of calcium transients being related to APs. But, clearly, the focus on high specificity led to false negatives as well (Fig. 1 B). Keeping the specificity at 100 % the average sensitivity of our transient detection was 73.3 ± 5.3 % (Fig. 1 C). Of note, a detection of the optical correlate of single APs was not achieved, employing this rigorous analysis aimed for maximizing specificity, even though we used state-of-the-art optics and indicators with high expression levels. The likelihood to identify an AP-related calcium transient increased with the number of APs exhibited by the GCaMP6f-positive neuron (Fig. 1 D) resulting in converging curves for AP and transient detection at higher firing rates (Fig. 1 E). In conclusion, while achieving high specificity comes at a cost, we could still capture 61.6 % of all electrophysiologically detected APs.

**Fig. 1.**
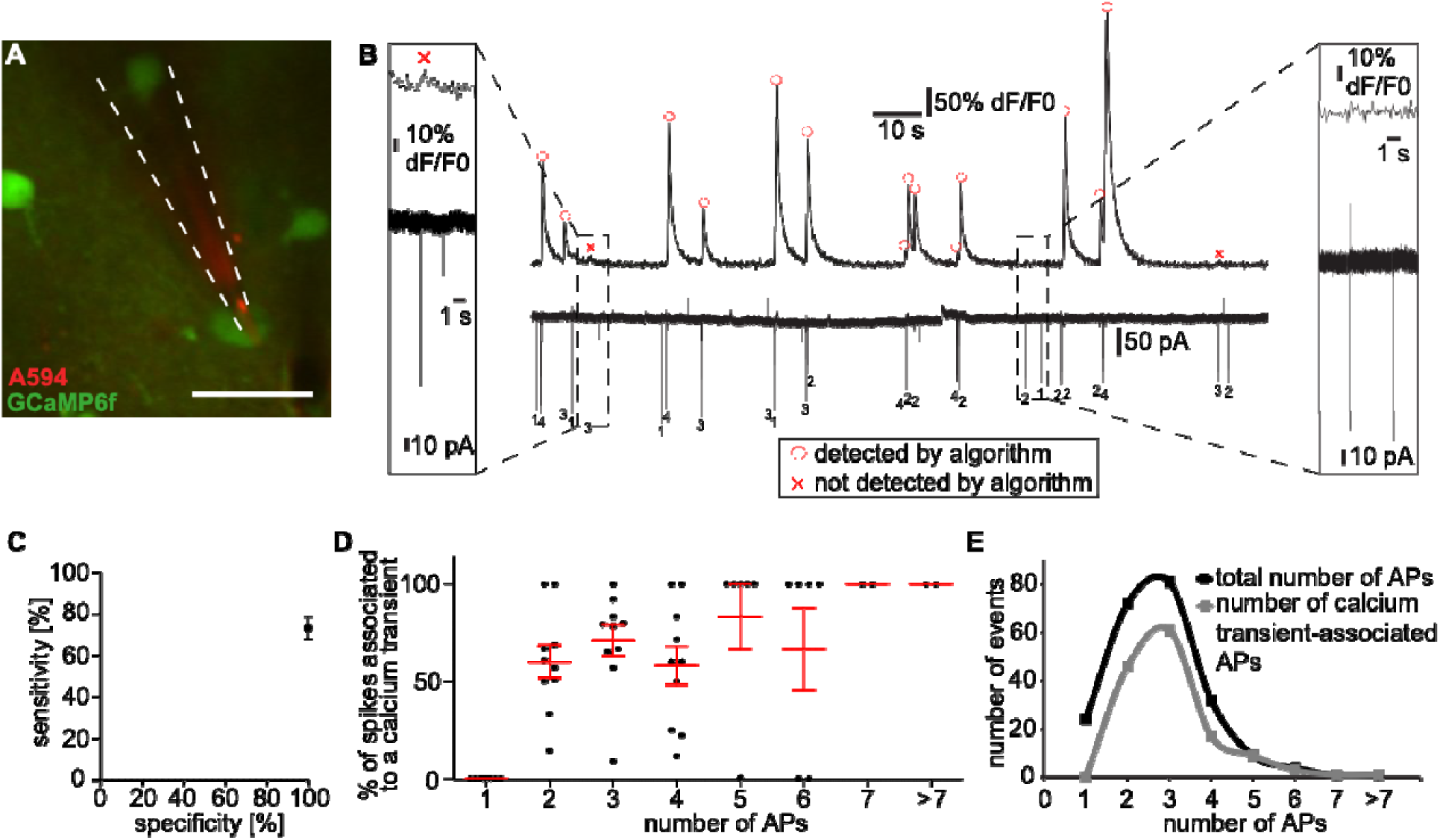
Implementing a highly specific optical detection of neural spiking activity in cortical circuits. **A** Simultaneous 2-P imaging and juxtacellular recordings of GCaMP6f-expressing neuron (green). Patch pipette filled with AlexaFluo594 (red), scale bar 20 µm. **B** Spontaneous activity of a GCaMP6f-expressing neuron. Calcium trace acquired with 2-P imaging (upper) and juxtacellular recording (lower). Detected calcium transient are indicated by a red circle. Undetected calcium transients are indicated by a red cross. Numbers of APs are indicated below the trace. Left magnified excerpt: Example of an AP-related calcium transient which has not been detected by the detection algorithm. Right magnified excerpt: Two AP events which do not have correlates in the corresponding calcium recording. **C** Sensitivity / specificity-plot, fixing specificity at 100 % results in an average sensitivity of 73.3 %, n = 7 neurons, 3 mice. **D** GCaMP6f sensitivity: Relation between spikes associated to a calcium transient and number of underlying APs. Mann-Whitney test 1 AP vs 2 AP p = 0.0012, 1 AP vs 3 AP p = 0.0012, 1 AP vs 4 AP p = 0.0012, Unpaired t-test 2 AP vs 3 AP p = 0.1521, 2 AP vs 4 AP p = 0.8938, 3 AP vs 4 AP p = 0.3628, n = 7 neurons, 3 mice. **E** Relation between number of underlying APs and number of events divided in total number of APs (black) and number of transient-associated APs (grey). Total number of APs, unpaired t-test 1 AP vs 2 AP p = 0.0627, 1 AP vs 3 AP p = 0.0311, 1 AP vs 4 AP p = 0.6729, 2 AP vs 3 AP p = 0.7439, 2 AP vs 4 AP p = 0.0709, 3 AP vs 4 AP p = 0.0331. Number of transient-associated APs, Mann-Whitney test 1 AP vs 2AP p = 0.0012, 1 AP vs 3 AP p = 0.0006, 1 AP vs 4 AP p = 0.0006, Unpaired t-test 2 AP vs 3 AP p = 0.4172, 2 AP vs 4 AP p = 0.0434, 3 AP vs 4 AP p = 0.0067, n = 7 neurons, 3 mice.

### Functional calcium transients can only be detected in a rather narrow spectral window

For an all-optical experiment with minimal cross-talk, it is essential to spectrally separate imaging and stimulation. Therefore, we needed to define the spectral window allowing for the reliable detection of functional calcium transients. We performed *in vivo* imaging of GCaMP6f-expressing neurons at different wavelengths of the same field of view, again in layer II/III of visual cortex in lightly anesthetized mice (Fig. 2 A). First, we investigated the shortest wavelength allowing for the identification of individual GCaMP6f-expressing neurons. The number of visually detectable neurons was similar across wavelengths ranging from 860 to 920 nm (Fig. 2 A). 860 nm was still in the pumping spectrum of the optical parametric oscillator (OPO, see below) and therefore might allow a system based on one laser source. However, while the neurons could be clearly identified, below 880 nm hardly any functional transient was detectable (18.3 ± 3.6 %, Fig. 2 B). At 880 nm a minor fraction (27.6 ± 10.0 %) and at 900 nm less than half of the transients could be recorded (48.8 ± 10.6 %) compared to imaging at 920 nm, where the number of detected calcium transients increased substantially (Fig. 2 B, D). Likewise, the number of detectable active cells increased in similar fashion when increasing the imaging wavelength from 860 to 920 nm (18.29 ± 3.59 % at 860 nm, 27.57 ± 9.93 % at 880 nm, 48.81 ± 10.57 % at 900 nm, 100 ± 0 % at 920nm, Fig. 2 C). Also, spontaneous transient frequencies increased significantly from 860 to 920 nm (0.33 ± 0.1 transients/min at 860 nm, 1.17 ± 0.29 transients/min at 880 nm, 1.33 ± 0.23 transients/min at 900 nm, 1.69 ± 0.22 transients/min at 920 nm, Fig. 2 E). Consequently, the spectral window for the detection of functional calcium transients is significantly smaller than the window for the morphological identification of GCaMP6f-expressing neurons.

**Fig. 2.**
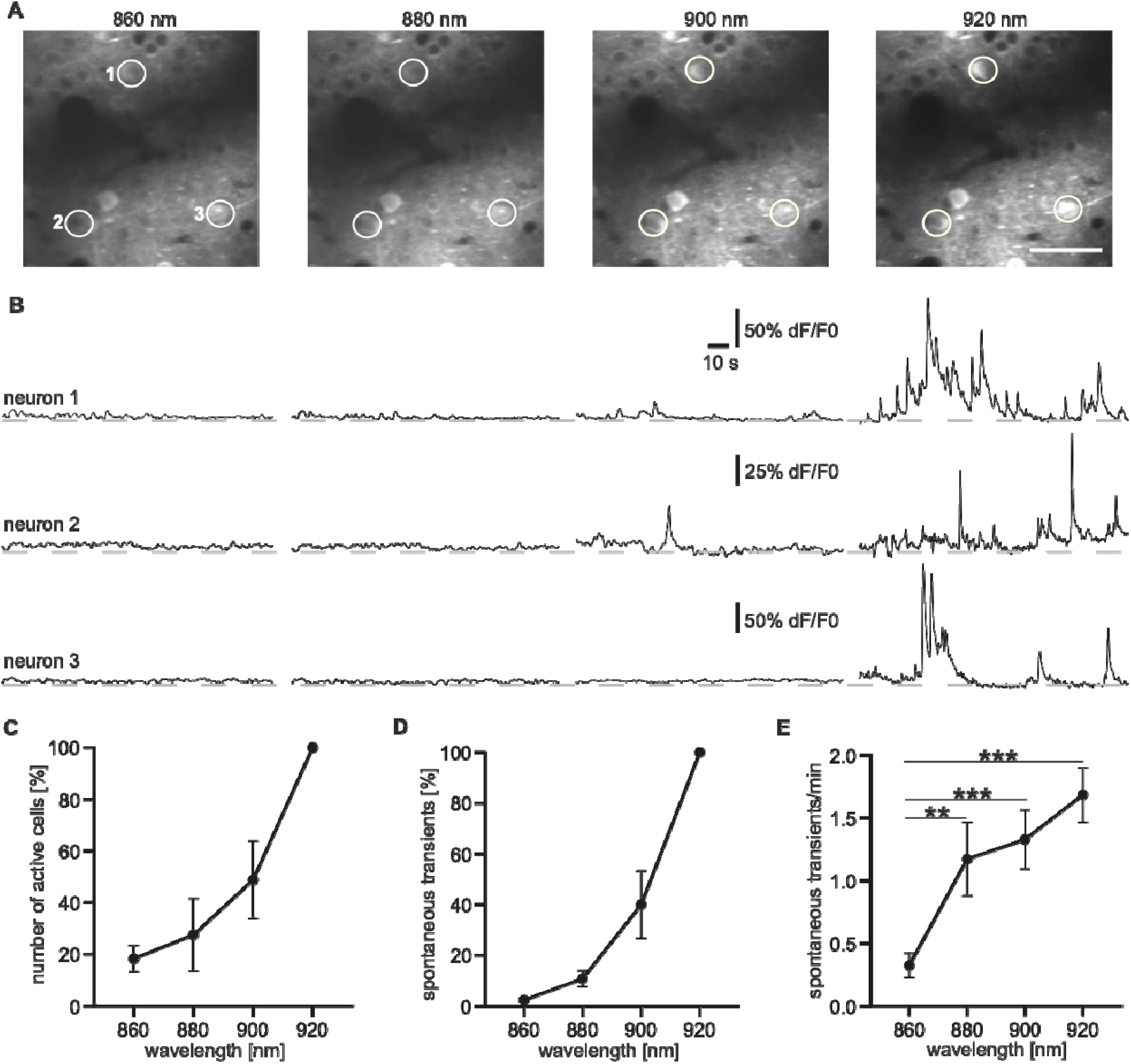
Functional calcium transients can only be detected in a rather narrow spectral window. **A** 2-P imaging of GCaMP6f expressing neurons at four different Ti:Sa wavelengths (860, 880, 900, 920 nm) in layer II/III of mouse visual cortex. The very same three neurons (white circles) are depicted at every wavelength. Scale bar 50 µm. **B** Corresponding calcium traces of depicted neurons. **C** Normalized number of active cells at different wavelengths. **D** Normalized number of calcium transients at tested wavelengths. **E** Average transient frequencies at different wavelengths. Mann-Whitney-Test, 860 nm vs. 880 nm p = 0.0026, 860 nm vs 900 nm p = 0.0003, 860 nm vs 920 nm, p = 0.0001, 880 nm vs. 900 nm p = 0.5069, 880 nm vs 920 nm p = 0.2800, 900 nm vs 920 nm p = 0.3757, n = 66 cells, 2 mice.

### Implementations of spectrally independent 2-P light sources

As the lower edge of the spectral range for our all-optical experiments was now set to 920 nm due to the functional limitations of GCaMP6f discussed above, we had to devise a light source, which flexibly delivered longer wavelengths for cross-talk free optogenetic modulations. Previous work on all-optical physiology used fixed-wavelength Ytterbium lasers typically between 1040 and 1080 nm for the 2-P excitation of opsins. We chose to probe whether longer wavelength might both reduce cross-talk and improve efficacy.

For achieving this goal, several technical solutions are amenable: One conceivable approach is to use one tunable 2-P Ti:Sa laser source and splitting the beam power for pumping an optical parametric oscillator (OPO) and for imaging GCaMP6f (Fig. 3 A). However, 920 nm, the wavelength for functional GCaMP6f imaging, is well-beyond the pumping spectrum of most OPO models. In this configuration the system could therefore only be used in combination with indicators effectively excitable with lower wavelengths, such as Oregon-Green-BAPTA 1 (OGB-1), excitable at 800 nm. For all-optical experiments reported in this study, based on GCaMP6f, we probed two different configurations both based on two primary laser sources: The first configuration integrates a second 2-P tunable Ti:Sa laser. One of the two independently tunable beams is dedicated for imaging, allowing full Ti:Sa-spectrum (680 - 1060 nm), and the other Ti:Sa is set at the optimal wavelength for pumping the subsequent OPO, resulting in full OPO-spectrum (1100 - 1400 nm) for optogenetic stimulation (Fig. 3 B). The second configuration comprises a single femtosecond laser with two independently tunable output channels simultaneously emitting two beams for imaging (680 - 960 nm) and stimulation (950 - 1300 nm) (Fig. 3 C). Each channel’s maximum output power exceeds 1 W. In both configurations, we temporally uncoupled dwell times of the excitation and imaging beams using separate lasers and scanners for imaging and stimulation. This is important, as the pixel dwell time achieved in resonant scanning mode is in the nanosecond range and therefore not sufficient for effective excitation of an opsin, mandating more than 4 µs *[26]*. Consequently, only the 2-P beam used for imaging was coupled to the fast resonant scanner and the second longer wavelength was coupled to an additional galvanometric-scanner. To achieve a high spatial specificity in modulating activity of single cells, it is crucial that the excitation volume is confined and does not exceed the spatial extent of the neuron supposed to be stimulated. Only then, specific scanning patterns can be applied to e.g. specifically target neurons showing aberrant activity without interfering the activity of well-functioning neighboring network components. The excitation volume of individual microscope systems is defined by their point spread function (PSF) and 2-P illumination theoretically allows excitation of femtoliter volumes. At 920 nm the PSF of our system had an acceptable full width half maximum (FWHM) of 0.20 ± 0.03 µm in x and 0.26 ± 0.01 µm in y direction, while the FWHM in z-direction was approximately 1.60 ± 0.05 µm (Fig. S1). These measurements where stable using different excitation wavelengths (Fig. S1) and allowed to concentrate the excitation on a 0.08 µm^3^ volume. In these configurations our 2-P system was therefore capable of performing spatially-specific and spectrally-independent optogenetic modulations temporally uncoupled from high-speed GCaMP6f-imaging.

**Fig. 3.**
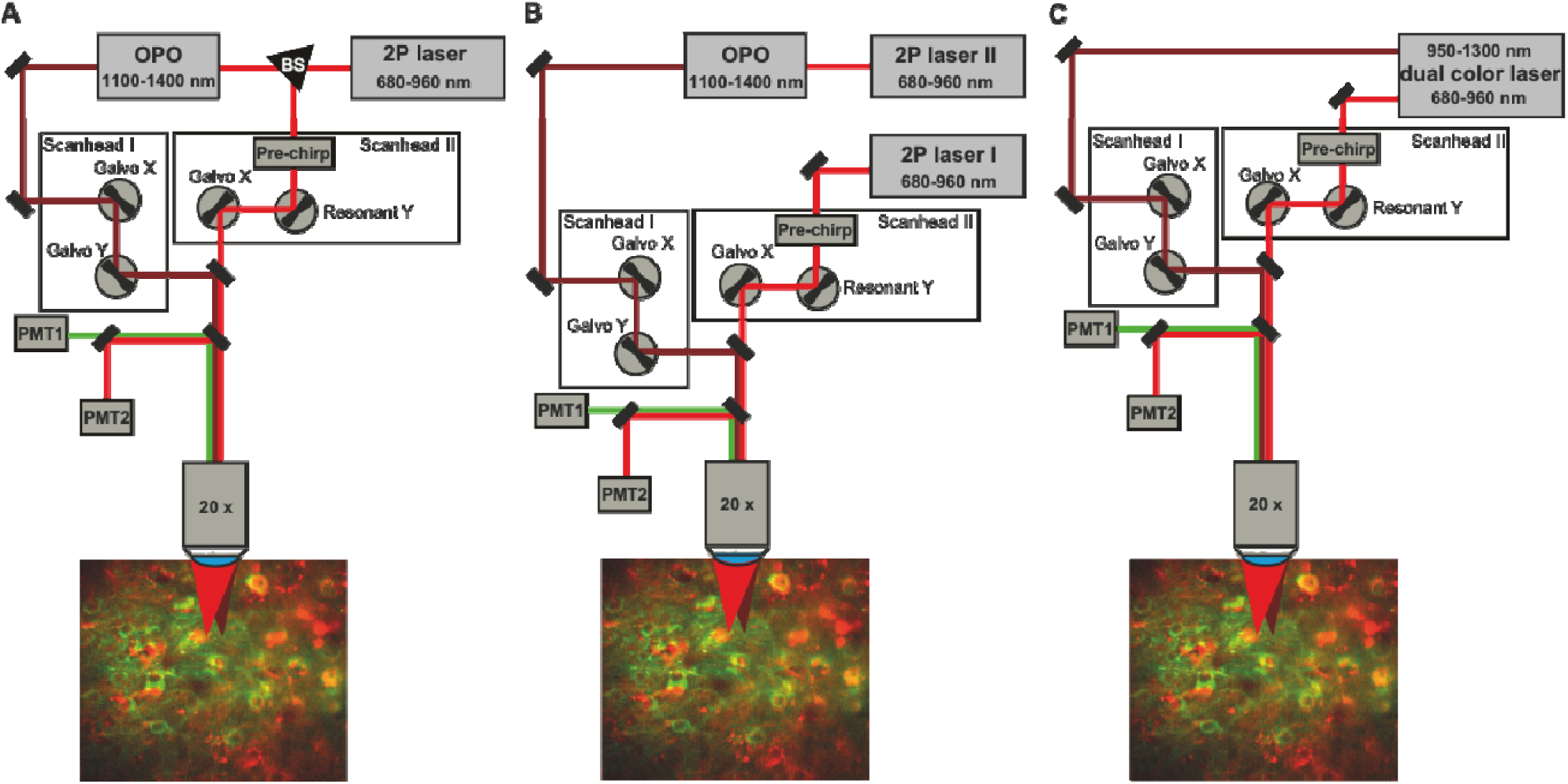
Extended-wavelength-spectrum 2-P all-optical interrogations can be achieved by using different light sources and system configurations. Schemes of a custom made 2-P microscope using different configurations and light sources. **A** Microscope set up based on one Ti:Sa laser used for both pumping the OPO and for GCaMP6f imaging by splitting the beam. The OPO delivers a broader wavelength range for excitation (1100 - 1400 nm). The imaging wavelength is guided to a resonant scanner for full-field imaging of GCaMP6f at 920 nm and the stimulation wavelength is guided to a temporally uncoupled galvo scanner for independent optogenetic control. PMT = Photomultiplier Tube, BS = Beamsplitter. **B** The same microscope configuration as in **A** is based on two independent Ti:Sa lasers, one is dedicated to GCaMP6f-imaging and the other one is pumping the OPO. **C** A dual color laser is delivering two independently tunable laser beams. Imaging wavelengths (680 - 960 nm) are guided to a resonant scanner and stimulation wavelengths (950 - 1300 nm) are guided to a separate galvo scanner a before.

### Careful titration of viral vectors is needed to achieve strong co-expression of both indicator and opsin

For highly specific all-optical interrogations combining precise optogenetic control and optical readout from individual neurons, the indicator and the opsin protein need to be co-expressed in the neuronal population of interest. We therefore co-expressed the 2-P excitable opsin C1V1_T/T_ in cortical excitatory neurons by performing AAV-based viral gene transfer using the excitatory neuron-specific promoter CaMKII together with GCaMP6f. The latter was expressed under the control of the ubiquitous hSyn-promoter and thereby gaining readout from all types of neurons. We titrated both viruses to achieve strong co-expression of both sensor and actuator protein by examining different dilutions of each virus. When increasing the titer of either AAV, we could observe a predominant expression of either GCaMP6f or C1V1_T/T_ -mCherry due to the limited protein synthesis capacity of single neurons (Fig. 4 A-C). Stable co-expression of C1V1_T/T_ and GCaMP6f could be observed in confocal analysis of the tissue sections employing a final titer of 20 : 1 for C1V1_T/T_ (3.33 * 10^11^ / ml) and GCaMP6f (1.84 * 10^10^ / ml) (Fig. 4 C). Using this titer the density of C1V1_T/T_-mCherry positive cells was 1392 ± 97 cells / mm^3^ in layer II/III and 4336 ± 313 cells / mm^3^ in layer V/VI (Fig. 4 D). The density of cells co-expressing C1V1_T/T_-mCherry and GCaMP6f was 735 ± 99 cells / mm^3^ in layer II/III and 1226 ± 139 cells / mm^3^ in layer V/VI (Fig. 4 E) resulting in a fraction of 52.5 ± 4.4 % and 28.3 ± 2.3 % of C1V1_T/T_/GCaMP6f co-expressing cells in layer II/III and in layer V/VI, respectively (Fig. 4 F). Hardly any co-expressing neurons were found in layer IV. We found strong and dense co-expression across targeted cortical layers II/III and V/VI. Note, that layer IV seems to be difficult to target with AAVs of serotype I/II, as reported previously *[28, 29]*.

**Fig. 4.**
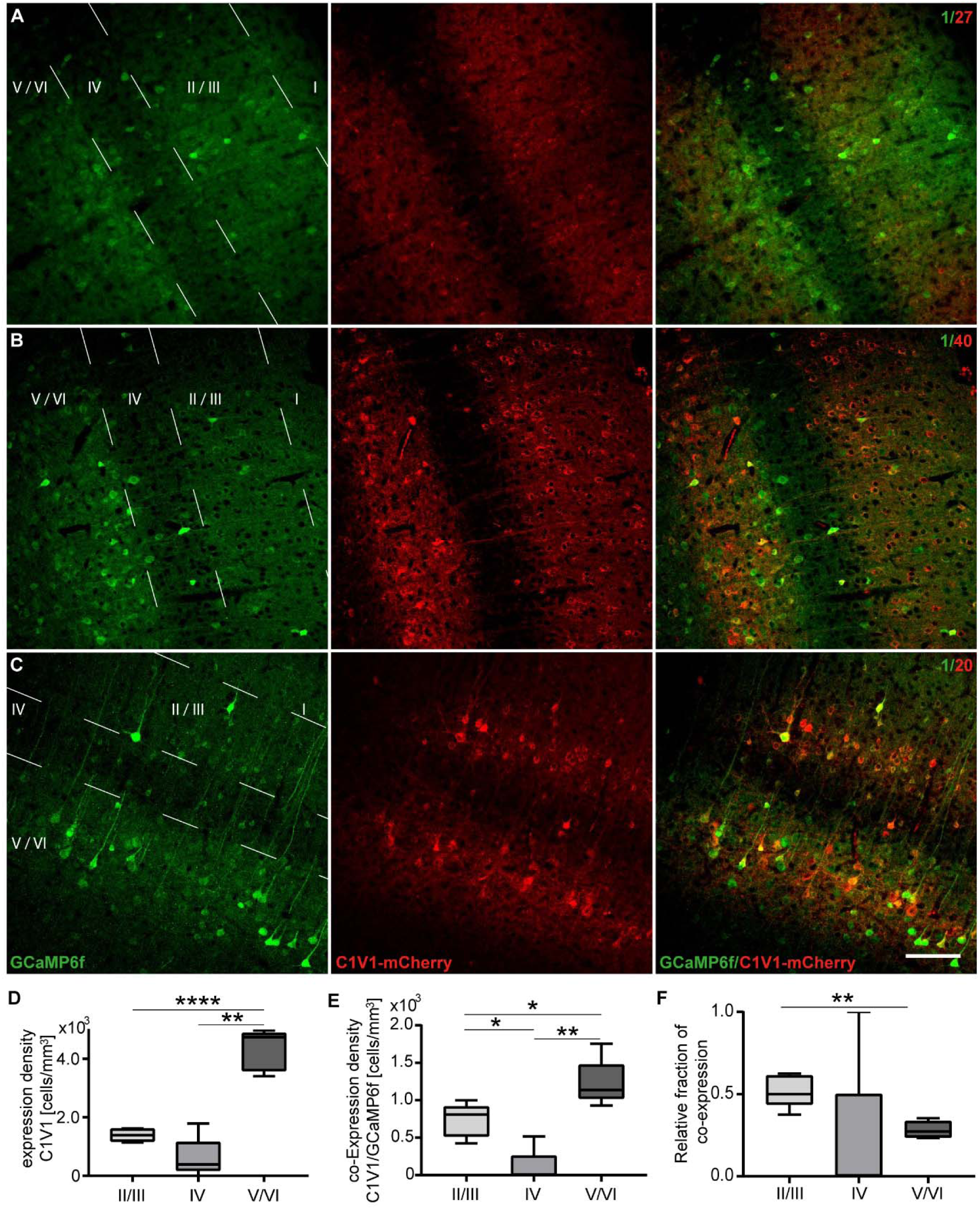
Strong co-expression of indicator and opsin. **A-C** Titration of AAVs encoding for indicator and opsin proteins: confocal micrographs of different titer combinations, dilution with PBS (GCaMP6f (green) / C1V1_T/T_ (red): [0.62 * 10^10^ / ml] / [6.66 * 10^11^ / ml], [0.92 * 10^10^ / ml] / [3.33 * 10^11^ / ml] and [1.84 * 10^10^ / ml] / [3.33 * 10^11^ / ml], 20x, scale bar 75 µm). Left column: green channel at 488 nm to evaluate GCaMP6f expression. Middle column: red channel at 584 nm to evaluate C1V1_T/T_-mCherry expression. Right column: merge of red and green channel to evaluate co-expression. **D** Density of C1V1_T/T_ expressing cells in different cortical layers of V1. Unpaired t-test, layer II/III vs layer V/VI p < 0.0001, Mann-Whitney test layer II/III vs layer IV p = 0.1508, layer IV vs layer V/VI p = 0.0079. **E** Density of C1V1_T/T_ / GCaMP6f co-expressing cells in different cortical layers of V1. Unpaired t-test layer II/III vs layer V/VI p = 0.0207, Mann-Whitney test layer II/III vs layer IV p = 0.0159, layer IV vs layer V/VI p = 0.0079. **F** Relative fraction of co-expressing cells in different cortical layers of V1. Unpaired t-test layer II/III vs layer V/VI p = 0.0014, Mann-Whitney test layer II/III vs layer IV p = 0.1111, layer IV vs layer V/VI p = 0.1111. **D-F** n = 175 cells, 5 brain slices (70 µm), 3 mice.

### Testing functional expression of the actuator C1V1 applying 1- and 2-P optogenetic stimulation

Electrophysiological recordings provide a direct readout of neuronal activity and are therefore well-suited to probe functionality of opsin expression and test 2-P excitation paradigms. To ensure functional expression of our optogenetic actuator, we therefore performed easy to implement optic fiber-based 1-P stimulations at 552 nm in combination with LFP-recordings (*[30]*, Fig. 5 A, B). Within single animals, we found a stable response pattern with a low variability of response amplitudes at approximately 60 mW / mm^2^ (Fig. 5 C). However, between animals amplitudes varied, with an average amplitude of 11.9 ± 0.4 mV. Next, we tested, whether our 2-P light source in combination with a raster scan paradigm was capable of eliciting spikes in individual opsin expressing neurons (Fig. 5 D). Simultaneous juxtacellular recordings demonstrated stable responses upon every trial of single-cell 2-P optogenetic stimulations with a light intensity of 37 mW at 1100 nm (Fig. 5 D, F). We applied a pixel dwell time of 6 µs with a raster width of 0.5 µm. Like this, specific modulations of single neurons were achievable by applying our stimulation approach with a success rate of 100 % using light intensities ≥ 37 mW in 26 consecutive trials (96 % at 18 mW). Latencies decreased with increasing light intensities (Fig. 5 G), as reported previously *[31]*. Interestingly, light intensities ≤ 37 mW only triggered approximately one event whereas light intensities around 60 mW already triggered two events (Fig. 5 H). This is important to note as for low AP numbers the sensitivity of our peak detection algorithm is decreasing substantially. Thus, we decided to perform our stimulations in imaging experiments at light intensities ≥ 40 mW. In summary, these quality-control and feasibility experiments provide the needed foundation for subsequent all-optical experiments.

**Fig. 5.**
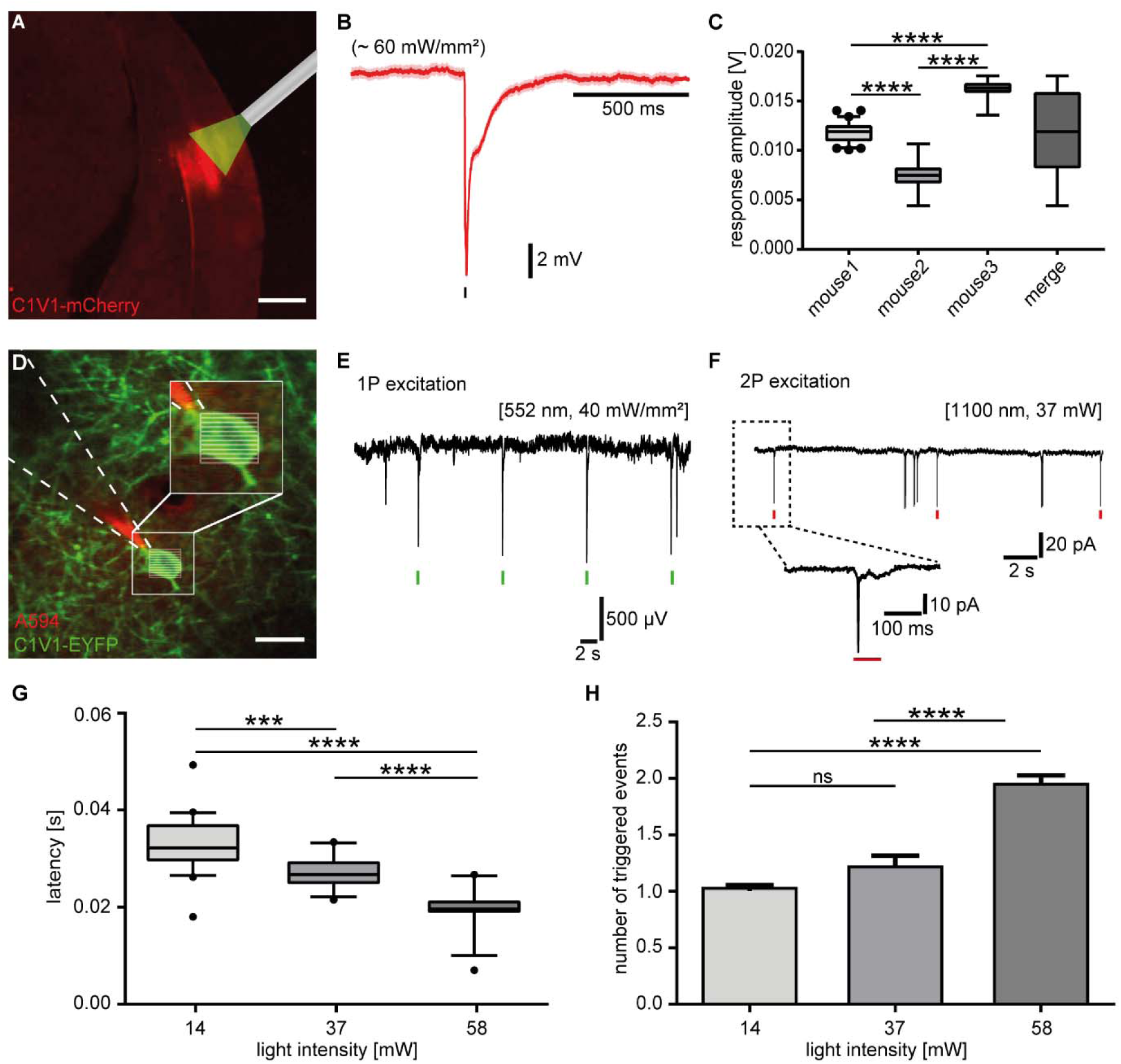
Probing functional expression of the actuator C1V1 applying 1-P and 2-P optogenetic stimulation. **A** Epifluorescent micrograph (5 x) of typical C1V1_T/T_-mCherry expression and schematic inset of the optic fiber for 1-P stimulation at 552 nm. Scale bar 500 µm. **B** Average LFP response upon optogenetic stimulation at 60 mW / mm2, n = 31 responses. **C** Average LFP response amplitudes across different animals and merged upon stimulations at 60 mW / mm2, unpaired t-test mouse 1 vs mouse 2 p < 0.0001, Mann-Whitney-Test mouse 1 vs mouse 3 p < 0.0001, mouse 2 vs mouse 3 p < 0.0001, n = 31 responses each, 3 mice. **D** *In vivo* 2-P image of shadow-patched C1V1-EYFP-expressing (green) neuron in layer II/III of mouse visual cortex and OPO-based 2-P stimulation using raster scan patterns at 1100 nm. Stimulation was delivered every 10 s and was defined by a duration of 68 ms, a pixel dwell time of 6 μs, a line scan resolution of 0.5 μm and a laser intensity of 37 mW. Patch pipette filled with AlexaFluo594 (red). Scale bar 15 µm. **E+F** Combined LFP and juxtacellular recordings revealed stable LFP responses upon 1-P stimulations at 552 nm (**E**) and robust action currents upon 2-P raster scans at 1100 nm (**F**), respectively. **G** Average latencies upon 2-P raster scan stimulations at different light intensities. Unpaired t-test, 14 mW vs 37 mW p = 0.0001, 14 mW vs 58 mW p < 0.0001, 37 mW vs 58 mW p < 0.0001, n = 26 responses, 1 neuron. **H** Average number of events triggered by 2-P raster scan stimulations at different light intensities. Mann-Whitney test, 14 mW vs 37 mW p = 0.06, 14 mW vs 58 mW p < 0.0001, 37 mW vs 58 mW p < 0.0001, n = 26 responses, 1 neuron.

### Above-noise stimulation artifacts of 2-P excitation are wavelength-dependent and absent above 1100 nm

Unlike electrophysiological recordings, optical imaging using genetically-encoded calcium indicators does not provide a specific signal per se. Identification of AP-derived calcium transients critically depends on several components of the signal, such as the sharp rise time. An artifact of the stimulation pulse in the GCaMP6f emission channel renders a highly specific identification of AP-related calcium transients difficult. This is of particular concern, if this method is applied in preclinical research, which should provide evidence guiding translation to human studies *[20-23]*. Here, we defined the spectral windows for artifact-free all-optical interrogations by testing different wavelengths for stimulation. To ensure that photostimulation artifacts were in fact no mistaken calcium transients, we performed our stimulation paradigm on mice which were lacking C1V1 expression and solely expressed GCaMP6f (Fig. 6 A). Notably, applying stimulation wavelengths and power used for all-optical experiments *[3, 5, 10, 12]*, we observed a significant artifact in the calcium trace when stimulating at 1020 nm, already at 40 mW power (Fig. 6 B). Increasing power levels to 80 mW which increases the efficacy of stimulation led to even more pronounced artifacts (Fig. 6 B, C). Note, that this artifact displayed a sharp rise time, and an amplitude well-above noise level, so that it might be misinterpreted as an AP-related calcium transient. Also, the amplitude of the artifact varied drastically between trials, rendering automated subtraction of mean artifacts, as conducted in previous study using single-wavelength all-optical physiology, obsolete *[28]*. To define the spectral range for artifact-free all-optical physiology, we now increased the stimulation wavelength step by step. We found that the amplitude of the averaged and normalized artifact decreased with increasing wavelength (Fig. 6 D). At 1060 nm, again, often used in Ytterbium-laser-based 2-P optogenetics, an artifact was still clearly visible, at least in one animal, at 80 mW in 35 of 70 analyzed cells (Fig. 6 D, E). At 1100 nm, no above-noise-level artifact could be observed (Fig. 6 D, E) except in one highly expressing cell (Fig. S2) of > 100 analyzed cells. In summary, it has to be stated that the artifacts induced by photostimulation were highly heterogeneous in shape and amplitude. This might be explained by unintended excitation of GCaMP’s fluorescent core eGFP and varies with varying expression levels of the reporter and varying levels of autofluorescence. We suggest using wavelengths of 1100 nm or higher for artifact-free all-optical physiology.

**Fig. 6.**
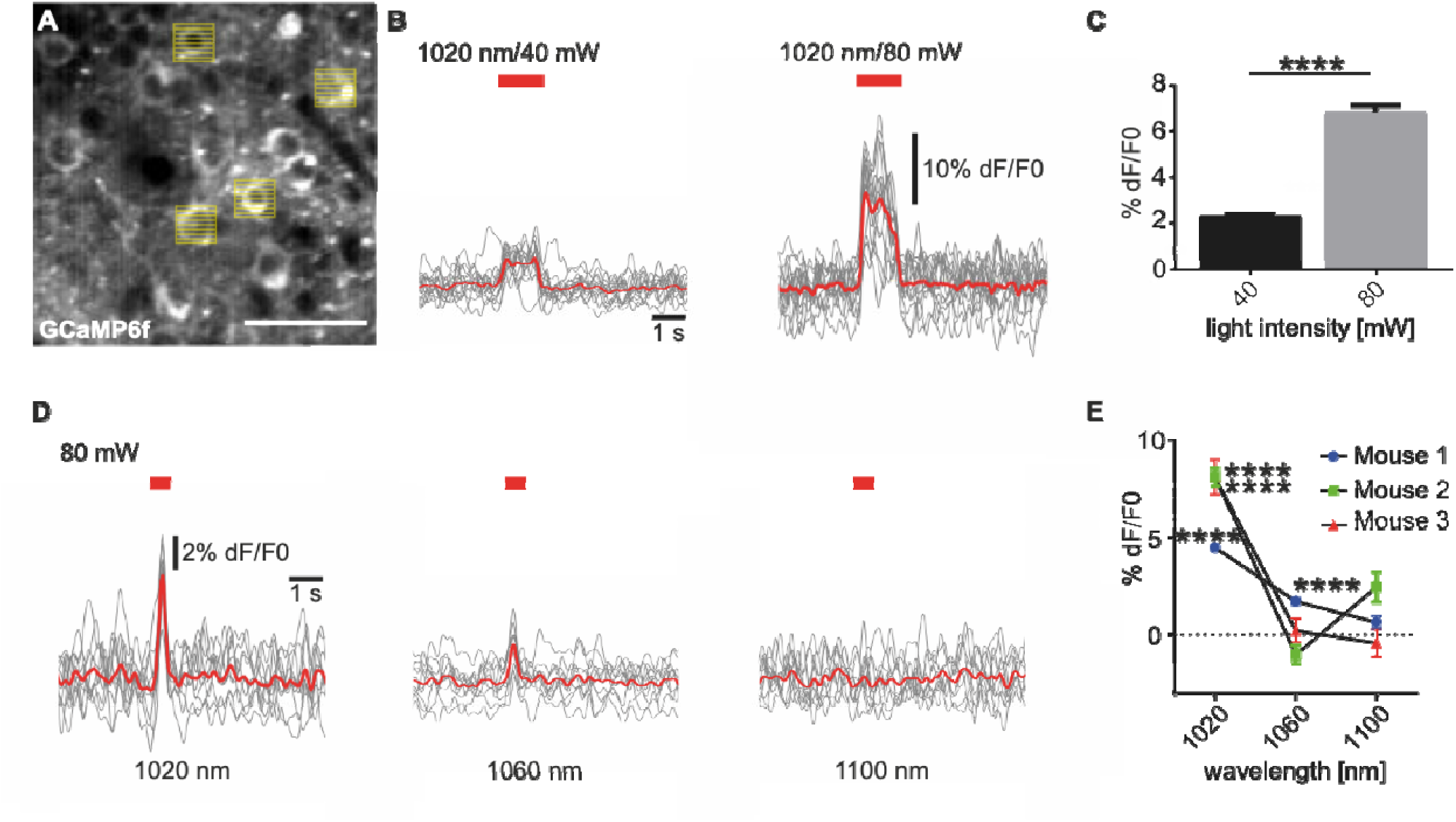
Stimulation artifact of 2-P excitation is wavelength-dependent and absent above 1100 nm. **A** *In vivo* 2-P calcium imaging of GCaMP6f expressing neurons in layer II/III of mouse visual cortex and simultaneous raster scan stimulations of selected cells at different wavelengths and light intensities. Please note the photostimulation artifact (white vertical lines). Scale bar 50 µm. **B** Average artifacts (red line, n = 10 artifacts, single trials are depicted in grey) in calcium traces of selected neurons upon 40 and 80 mW raster scan stimulations at 1020 nm. The amplitude of the artifact is increasing with increasing stimulation light power. **C** Quantification of artifact amplitude at 40 and 80 mW. Mann-Whitney test 40 mW vs 80 mW p < 0.0001. 40 mW n = 1192 trials, 2 mice, 80 mW n = 1520 trials, 3 mice. **D** Averaged artifact (red lines, n = 10 artifacts, single trials are depicted in grey) upon 2-P raster scan stimulations (80 mW) at varying stimulation wavelengths (1020, 1060 and 1100 nm). **E** Quantification of artifact amplitude at different stimulation (80 mW) wavelengths. Average artifact amplitudes are decreasing with increasing wavelength. Please note, that at 1100 nm no above-noise artifact is observable. In all mice and conditions mean artifacts were tested vs. standard deviation of baseline level, Mann-Whitney test, Mouse 1 1020 nm, n = 530 trials, 53 neurons, p < 0.0001, 1060 nm, n = 700 trials, 70 neurons, p < 0.001, 1100 nm, n = 250 trials, 25 neurons, p = 0.0223, Mouse 2, 1020 nm, n = 540 trials, 54 neurons, p < 0.0001, 1060 nm, n = 690 trials, 69 neurons, p < 0.0001, 1100 nm, n = 690, 69 neurons, p = 0.1617, Mouse 3 1020 nm, n = 450, 45 neurons, p < 0.0001, 1060 nm, n = 195 trials, 15 neurons, p = 0.6236, 1100 nm, n = 360 trials, 36 neurons, p < 0.0001.

### 1100 nm is the optimal wavelength for effective and cross-talk-free optogenetic control

Finally, we conducted *in vivo* experiments in animals expressing high and functional levels of both opsin and indicator in the lightly anesthetized mouse. First, we performed GCaMP6f imaging only. We assessed simultaneously the full-field activity of the local ensemble of neurons. We set the anesthetic depth in a way that persistent, desynchronized population activity could be observed (Fig. S3). Deeper anesthesia will result in slow-wave-brain state, which is characterized by large-amplitude oscillations (Fig. S3). This brain state would not be suitable for probing the excitability and connectivity of each neuron, as the excitability of the respective neuron would be governed by the current phase of the population-wide oscillation *[32]*. In persistent brain state, we next carried out cell-specific interventions on genetically-defined C1V1_T/T_-expressing neurons using 2-P based optogenetic stimulation. We carried out sequential stimulations of individual cells at the artifact-free wavelength of 1100 nm using aforementioned raster scans while simultaneously imaging GCaMP6f-fluorescence at 920 nm (Fig. 7 A). The performed stimulation paradigm robustly elicited putatively AP-related GCaMP6f transients in stimulated cells, applying the highly specific detection algorithm implemented above (Fig. 7 B). We now asked, whether increasing stimulation wavelengths beyond 1100 nm, not being possible with standard Ti:Sa or Ytterbium lasers, will lead to an increased efficacy. As stated above, the early studies in the field revealed a linear increase in photocurrents, until the technical limit of 1040 nm *[26]*. It has to be noted, however, that these studies were conducted in slice preparations, with a drastically different inhibitory tone. As the main goal of this methodological pipeline is testing the limitations and prospects of 2-P all-optical physiology in preclinical *in vivo* applications, we believe, that measuring the capability of evoking detectable and specific AP-associated calcium transients represent the most appropriate avenue. Surprisingly, we found that a further increase of the stimulation wavelength from 1100 up to 1300 nm did not yield a significant increase in the rate of evoked calcium transients (Fig. 7 C) as well as no significant increase in the fraction of responding cells (Fig. 7 C). However, increasing stimulation power from rather low values ranging at 40 or 80 mW up to values ranging at 120 or 210 mW significantly increased the rate of evoked transient 2-3 fold (Fig. 7 D, 26.7 ± 6.7 % at 40 mW, 21.6 ± 3.5 % at 80 mW, 56.7 ± 13.1 % at 120 mW, 76.3 ± 7.3 % at 210 mW), but, also here, increasing light intensities did not impact the fraction of responding cells (Fig. 7 D).

**Fig. 7.**
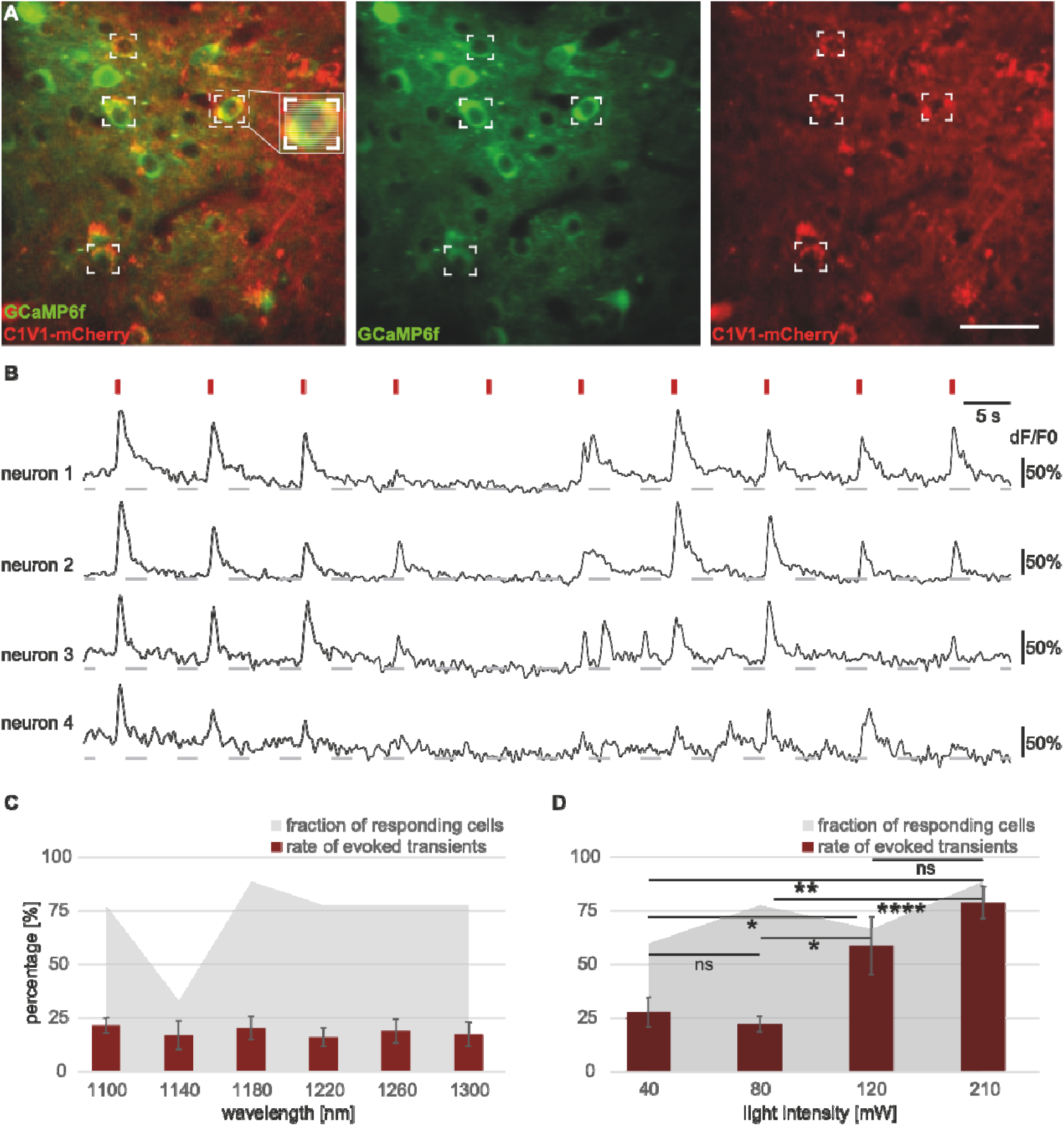
1100 nm represents the optimal wavelength for effective and cross-talk-free optogenetic control. **A** All-optical control of individual GCaMP6f (green) / C1V1_T/T_ (red) co-expressing neurons in layer II/III of mouse visual cortex. Depiction of raster scan patterns a described above. Six individual neurons can be targeted for sequential photostimulation. Scale bar 50 µm. **B** 2-P stimulation (indicated by red marker) of GCaMP6f / C1V1_T/T_ co-expressing neurons at 1100 nm. GCaMP6f calcium transients of four co-expressing neurons upon ten stimulation trials. **C** Average rate of evoked transients (red bars) at varying wavelengths (1100 - 1300 nm) at 80 mW. Grey shadow indicates the fraction of responding cells. n = 9 neurons, 5-10 trials each, 1 mouse. **D** Same as in **C** but varying light intensities (40, 80, 120 and 210 mW) at 1100 nm. Same color coding as in **C**, n = 23 neurons, 5-10 trials each, 3 mice. Unpaired t-test, 40 mW vs 80 mW p = 0.4570, 40 mW vs 120 mW p = 0.01227, 40 mW vs 210 mW p = 0.0039, 80 mW vs 120 mW p = 0.0115, 80 mW vs 210 mW p < 0.0001, 120 mW vs 210 mW p = 0.1886.

## Discussion

In this study, we provide a methodological and conceptional framework for a highly specific all-optical physiology approach geared at applications in network-based therapies for preclinical research. For that, challenges had to be overcome in devising a light source for spectrally independent imaging and optogenetics. Here, we used a dual-scanner approach, which is well-suited for establishing this framework. In our view, holographic or combined holographic and scanning approaches based on spatial light modulators *[3-5, 10, 12]*, temporal focusing *[2, 9]* or new techniques such as 3D-SHOT *[7]* are superior to scanning approaches as used in this study as they allow temporally uncoupled, parallel manipulation and imaging of a multitude of cells in three dimensions. However, holography-based systems are typically set at defined stimulation wavelengths as they are often used with fixed-wavelength Ytterbium lasers, which do not allow for flexible exploration of spectral windows for effective and artifact-free optogenetic control. Therefore, a 2-P scanner-based solution as applied here might still be advantageous at least for methodological ground work in the field of all-optical physiology. It has to be noted, that in terms of light sources, novel designs providing two independently tunable femtosecond-pulsed laser beams with extended wavelength spectrum, could be introduced in CGH-based setups as well, thereby increasing flexibility in contrast to fixed-wavelength lasers.

We commenced our framework with setting the specificity to detect underlying APs of our detection algorithm to 100 % by correlating our optical data to the actual electrical activity of a neuron. This allowed to completely avoid type I errors, i.e. mistaking movement- or stimulation light-induced artifacts with calcium transients devoid of an electrophysiological correlate and therefore contributing to false-positive detection of pathological activity patterns (e.g. early cortical hyperactivity) in a healthy brain. Here, preclinical research has to tackle the current reproducibility crisis: both in preclinical models of neurodegenerative disease such as Alzheimer’s or Huntington’s disease as well as models of cerebral stroke, decades of research devised a multitude of drugs showing highly promising effects in rodent models, however, not a single therapeutic concept proved effective in the human clinical setting. Surely, the reasons for that are diverse, but, the avoidance of false-positives has been identified as one major obstacle *[20-23]*. Therefore, the field has to rethink the entire workflow, comprising the experiments per se, but also the analysis of neuronal activity. In our recent studies, we already applied a rigorous analysis routine to 2-P imaging data. And indeed, this provided an improved platform for testing the efficacy of drug treatments *[14, 15]*. Here, we extend this greatly conservative approach to all-optical physiology. Of course, this comes at cost of sensitivity of activity detection, especially of low numbers of APs. It has to be noted, that the initial proof-of-principle publications on new indicators, e.g. on GCaMP6f *[33]* do provide sensitivity, but no specificity data. Certainly, also in our setup, we did identify calcium deflections upon single APs, but, these could not be unambiguously separated from calcium deflections of non-physiological sources. In addition, the specificity of these approaches is compromised by systematic artifacts induced by photostimulation. We here demonstrate that the amplitude of these artifacts can be in the range of the amplitude of calcium transients and define the spectral range for artifact-free all-optical experiments. On the basis of this finding, we cannot recommend automated analysis algorithms, which merely correlate normalized fluorescence values and strongly suggest that a detection algorithm should be based on an event detection, i.e. the optical correlate of a neuronal action potential. Indeed, several elegant approaches take these characteristic dynamics into account, such as the peeling algorithm *[34]*. But, ultimately, given the high variability of technical setups, expression levels and brain areas investigated, we still suggest employing single-cell electrophysiology to titrate each detection algorithm, as exemplified in this study. Of course, in 2-P all-optical experiments these artifacts are less dramatic than in 1-P excitation *[28]*. However, also in 2-P excitation these photostimulation artifacts still cause significant problems *[2, 4, 6, 7, 9, 10]* and might resemble AP-related calcium transients *[5]*, especially when high light intensities are applied. Up to now, this problem was addressed using three main approaches: 1.) Sacrificing pixels containing the artifact resulting in data loss *[9, 10]*, 2.) post-hoc identification based on the temporal dynamics of the artifact in contrast to the AP-related transients *[2, 4]*, which in our view is amenable only in data with highest signal-to-noise ratio, not reflecting the reality particularly in awake measurements and 3.) gating stimulation and recording lasers, reducing effective field of view *[7]*. In contrast to the aforementioned studies, one study used a different configuration combining a blue-shifted opsin with a red-shifted indicator *[6]*. However, also here, a photostimulation artifact was apparent, most likely due to the fact that the indicator was excited on his blue shoulder by the strong stimulation light, and was dealt with by subtracting background or blanking a short period in the calcium trace *[6]*. One further, major problem which cannot be addressed with any of these methods and is independent of the artifact amplitude: The photostimulation artifact will inevitably make it impossible to determine latencies of responses and will therefore hinder precise timing of evoked spikes as the exact onset of a transient will be hidden within the artifact.

Here, we provide evidence that at stimulation wavelengths of 1100 nm and above, no above-noise-level artifacts can be detected. This frees the researcher from the need for any post-hoc and specificity-reducing methods of artifact minimization and opens up the use of higher stimulation light intensities, e.g. to overcome strong inhibitory tones, as reducing light intensities to minimize artifacts will necessarily reduce photocurrents *[2, 7, 9, 10]*. Indeed, increasing stimulation light intensities at the artifact-free wavelength of 1100 nm significantly increased cellular response rates. And above all, stimulation at 1100 nm will still allow the exact determination of response latencies.

Finally, we addressed an open question of the field in terms of the spectral efficacy of opsin activation beyond 1100 nm. As aforementioned, *in vitro* whole-cell electrophysiology measurements found an increase in photocurrents with increasing wavelengths up to 1040 nm *[26]*. Here, albeit only using C1V1 in *in vivo* 2-P calcium imaging experiments, we did not find a further significant increase in spectral efficacy beyond 1100 nm. However, this might be different for other 2-P excitable opsins and, here, is an important limitation to our study: We are only probing this framework for a single indicator-opsin-pair, as this combination seemed to be mostly used in the field *[2-5, 8, 12]*. However, the development of new opsin and indicator variants has advanced *[7, 9, 10, 35]* and for newer opsin-indicator-couples different stimulation or imaging parameters might be optimal. Furthermore, as mentioned above our approach has important technical limitations compared to CGH-based all-optical experiments and is currently limited to one plane for imaging and stimulation.

In conclusion, the still new and evolving field of 2-P all-optical physiology opens up tremendous prospects, also in the field of preclinical research aiming for network-based therapies *[19, 36]*, but, we suggest a careful and holistic approach including a pipeline for further exploratory research as devised here in order to maximize translational power.

## Materials and Methods

### Animals

All experiments were carried out along institutional animal welfare guidelines and approved by the Landesuntersuchungsamt Koblenz, State of Rhineland-Palantine, Germany. For experiments adult female and male C57/BL6 mice were used.

### Study design

The overarching research goal was to improve all-optical approaches for their use in translational models of neurological disorders. Here we pursued three a priori defined research objectives: (I.) Achieving utmost specificity in terms of the detection of AP-related calcium transients, (II.) investigating the currently maximally achievable spectral range for all-optical experiments concerning, both, imaging and stimulation and to determine the interference of photostimulation artifacts, and (III.) probing the wavelength dependency of effective optogenetic control. The study is an explorative study on non-randomized female and male C57/BL6 mice co-expressing C1V1 and GCaMP6f in cortical neurons. It represents a completely new experimental paradigm with designated hardware components, which has not been used in this field of research, yet. Key designated components of this study are a state-of-the-art 2-P microscope integrated with an OPO or a Cronus laser, C1V1 as optogenetic actuator and GCaMP6f as calcium indicator. Particularly in experiments exploring the three a priori defined research objectives (Fig. 1, Fig. 6 and Fig. 7) sample sizes were determined by power analysis using the effect size and accuracy from our earlier studies. Light power levels for optogenetic stimulation were based on previous studies *[3, 26]*.

### Stereotactic virus injections and chronic window preparations

For 2-P all-optical experiments GCaMP6f was co-expressed with the opsin C1V1_T/T_. This was achieved by titration of both viral solutions. Stock solutions of AAV1.Syn.Flex.GCaMP6f-WPRE.SV40 (3.67 × 10^11^ / mL, Penn Vector Core, University of Pennsylvania, PA, USA) and rAAV2/CamKIIa-C1V1(E122T/E162T)-TS-mCherry (1.0 × 10^12^ / ml, UNC Vector Core, NC, USA) were prepared by diluting with PBS and stored at 4 °C. Different titers were tested ([0.62 × 10^10^ / ml] / [6.66 × 10^11^ / ml], [0.92 × 10^10^ / ml] / [3.33 × 10^11^ / ml] and [1.84 × 10^10^ / ml] / [3.33 × 10^11^ / ml]). For LFP experiments rAAV2/Ef1a-DIO-C1V1(E122T/E162T)-TS-mCherry (UNC Vector Core, NC, USA) was injected alongside with rAAV2/PKG-Cre (UNC Vector Core, NC, USA) in a ratio of [1.44 × 10^13^ / ml] / [4.0 × 10^12^ / ml]. Adult C57/BL6 mice anesthetized with isoflurane in O_2_ (Abbvie, Ludwigshafen, Deutschland) were placed in a stereotactic frame (Kopf, CA, USA) and warmed with a heating pad (World Precision Instruments, Sarasota, FL, USA). For virus injections, a craniotomy was carried out above V1 with 3.0 mm posterior to Bregma and 2.5 mm lateral to the midline. Viral constructs were delivered though a small durotomy by a glass pipette (Hirschmann Laborgeräte, Eberstadt, Deutschland) via a custom-made loading system including a syringe and a plastic tubing using manual pressure. The pipette was slowly inserted and approximately 300 nl of the viral solution was injected with an injection-speed of 0.1 µl / min *[37]* at a depth of 200 µm targeting layer II/III and 600 µm targeting layer V/VI. Before retraction, the pipette remained in place for 5 minutes preventing an outward flow of the viral solution. For chronic window preparations, the skull was exposed to fix the head-holder on the mouse’s head. Using the identical stereotactic coordinates as for histology a circular cranial window was prepared above V1. The virus injection procedure was conducted as described above. After the injection of the viral constructs, the opening was closed with a circular cover slip (Electron Microscopy Sciences, Hatfield, PA, USA) at a diameter of 4 mm.

### Histology, confocal imaging and quantification of co-expression

For characterization of co-expression, animals were perfused trancardially with 4 % PFA 5 weeks post injection. Brains were postfixed in 4 % PFA for 24 hours until they were sliced (70 µm thickness) using a Leica Vibratome (Leica, Wetzlar, Germany). Slices were directly mounted on an object slide (Gerhard Menzel, Braunschweig, Deutschland) with Vectashield Mounting Medium (Vector Laboratories, Burlingame, CA, USA) and fixed with a cover slip (Gerhard Menzel, Braunschweig, Deutschland). Brain tissue sections were analyzed with a Leica SP8 confocal microscope (Leica, Wetzlar, Germany) equipped with a Leica 20 x HCX PL APO dry (NA = 0.75) objective. To determine the density of C1V1_T/T_ expressing and C1V1_T/T_ / GCaMP6f co-expressing cells in V1, all cells with smooth and strong expression within the volume of strong homogeneous fluorescence were counted separately for each layer by using the cell counter plug-in of ImageJ (https://imagej.nih.gov/ij/). The density of cells per mm^3^ in cortical layers II/III, IV, V/VI was calculated by dividing the total number of cells within the volume of strong homogenous fluorescence in each layer by the volume. Expression densities and fraction of expressing cells of all counted slices were averaged for each layer by calculating arithmetic means.

### *In vivo* electrophysiology

For LFP recordings mice were injected with rAAV2/Efla-DIO-C1V1-(E122T/E162T)-TS-mCherry + rAAV2/PKG-Cre as described above. General anesthesia was performed with an intraperitoneal injection of 2.5 µl / g containing 0.5 µl Medetomidin (Pfizer AG, New York, USA; 1 mg / ml), 1 µl Midazolam (Roche Pharma AG, Basel, Switzerland; 5 mg / ml) and 1 µl Fentanyl (Sintetica S.A., Mendrisio, Switzerland; 50 µg / ml). The coordinates of preceding virus injection were set and an acute 2 × 2 mm cranial window was prepared using a dental drill. With a fluorescence lamp, the localization of the virus expressing cortical area was determined. For LFP recordings patch-pipettes (1.5 - 3 MΩ) were filled with PBS and inserted into the brain tissue close to the virus expressing region in a depth of - 300 µm. The LFP signals were recorded with an EXT-02F/2 amplifier (npi Electronic, Tamm, Germany). The signal was filtered low pass at 300 Hz without high pass filter. Optogenetic stimulation was delivered by a 20 mW solid-state laser at 552 nm coupled with an acoustic-optic modulator (Crystal Technology, Palo Alto, CA) for rapid control of laser intensity. The laser beam was bundled into an optic multimode fiber (Thorlabs, Grünberg, Germany) using a fiber collimator (Thorlabs, Grünberg, Germany). The fiber had an outer diameter of 0.2 mm, a core diameter of 0.1 mm and a NA of 0.48. The fiber was guided along the arm of the stereotactic setup and placed above the virus expressing region in physical contact with the intact dura mater. Single 10 ms stimuli were applied with approximately 60 mW / mm^2^ light density. The voltage signals were collected with an analog-digital converter (CED, Cambridge, UK), displayed in real-time and saved with the appertaining software Spike 2 (CED, Cambridge, UK) on a computer (Dell Inc., Round Rock, TX, USA). The data was collected with a recording frequency of 1 kHz. Amplitudes were analyzed with Igor Pro (WaveMetrics, Portland, OR, USA) and quantified with GraphPad (GraphPad Software Inc., La Jolla, CA, USA). Average response amplitudes were calculated by using the arithmetic mean.

For juxtacellular recordings mice were injected with AAV1-Syn-GCaMP6f-WPRE-SV40, AAV2-CaMKIIa-C1V1(E122T/E162T)-TS-mCherry or AAV2-CaMKIIa-C1V1(E122T/E162T)-TS-EYFP. Three weeks later, the mouse was anesthetized with isoflurane and an acute craniotomy above V1 was prepared as described above. The exposed cortex was superfused with warm PBS. To dampen heartbeat- and breathing-induced motion, the cranial window was filled with 1 % agarose. Juxtacellular recordings were made by visually targeting GCaMP6f- or C1V1-expressing cells under 2-P vision in layer II/III of V1. Patch electrodes (4 – 6 MΩ) were filled (in mM): 10 HEPES, 1 MgCl_2_, 2 CaCl_2_, 150 NaCl, 2.5 KCl, 20 glucose and 10 Alexa Fluor 594 or Alexa Fluro 488 for pipette visualization. Signals were recorded with an ELC-03XS amplifier (npi Electronic, Tamm, Germany). Squarish ROIs (∼ 30 × 30 µm) were placed onto the C1V1-expressing cells and raster scanning was performed. Stimulation was delivered every 10 s and was defined by a duration of 68 ms, a pixel dwell time of 6 µs and a line scan resolution of 0.5 µm. The maximum laser intensity applied was 147 mW at 1100 nm. The voltage signals were collected and analyzed as described above.

### 2-P calcium imaging and 2-P optogenetic stimulations

Chronic imaging experiments were performed earliest four weeks post injection to ensure sufficient co-expression of GCaMP6f and C1V1_T/T_. For imaging animals were anesthetized with isoflurane and fixed on a custom-made table. An isoflurane dose of 1.0 - 1.5 % isoflurane / O_2_ resulted in an average breathing rate of 50 - 70 bpm and reliably induced slow-oscillatory state with its characteristic Up/Down state transitions. An anesthesia dose of 0.6 - 1.0 % lead to recordings in persistent state. Here the breathing rate of the animal was 100 - 110 bpm. The custom-made 2-P microscope set-up (LaVision Biotec, Bielefeld, Germany) was equipped with a resonant scanner for fast full-field scanning up to 35 Hz. A mode-locked femtosecond-pulsed Tisa:Sapphire laser (Coherent, CA, USA) was used at 860, 880, 900 and 920 nm for imaging GCaMP6f expressing cells. The laser operated with 1-20 % of its maximal power output (3 W). The imaging plane was usually between 250 ± 100 µm below the cortical surface and the field of view was 466 × 466 µm using a Zeiss W-Plan-Apochromatic 20 x DIC VIS-IR objective (NA = 1.0). For all-optical experiments with GCaMP6f / C1V1_T/T_ two different technical approaches were used. In the first implementation, the imaging was performed with the aforementioned Ti:Sapphire laser and the 2-P excitation of C1V1_T/T_ was conducted with an optical parametric oscillator (Coherent, CA, USA) delivering light between with (1100 - 1400 nm) and pumped by a second independent Ti:Sa laser. Alternatively, both imaging and excitation were performed with a single femtosecond laser with two independently tunable output channels (Light Conversion Ltd., Vilnius, Lithuania). The first tunable channel (680 - 960 nm) was used for 2-P imaging, the second tunable channel (950 - 1300 nm) was used for 2-P excitation of C1V1_T/T_. The laser operated with 1-30 % of its maximal output power (1.2 W) in the imaging channel and 1-50 % of its maximal output power in the excitation channel (1.1 W). The pulse durations of both channels were 130 ± 20 fs across the wavelength tuning ranges, provided by manufacturer. The pulses were synchronous with repetition rate of 76 MHz each. In both cases only the 2-P beam used for imaging was coupled to the fast resonant scanner and the second longer wavelength was coupled to the additional galvanometric-scanner. Laser intensity was controlled by an electro-optical modulator (Coherent, CA, USA). Two 1-P lasers at 488 and 561 nm (High Performance OBIS_TM_ Laser Systems, Coherent, Santa Clara, CA) were also coupled to the additional galvanometric scanner. For signal collection a high sensitive PMT (Hamamatsu, Hamamatsu, Japan) was used. Multiple C1V1_T/T_ expressing neurons were stimulated individually or in full-field manner. Using ROI-based stimulation up to 6 ROIs were placed onto the cells of interest and raster scanning of each ROI was performed. Stimulation was delivered every 10 s and was defined by a pixel dwell time of 6 µs and a line scan resolution of 0.5 - 2 µm. The maximum laser intensity applied was 210 mW at 1100 nm. The nanobead sample (Invitrogen, Carlsbad, CA, USA) was used to measure point spread functions across wavelength tuning ranges for imaging and excitation. Images of fluorescent nanobeads and FWHM values were extracted in all three dimensions and fitted using a gaussian distribution.

### 2-P calcium imaging data analysis

The analysis of 2-P calcium imaging data was done stepwise. Firstly, the calcium imaging data was acquired with Imspector (LaVision Biotec, Bielefeld, Germany). Afterwards image series were imported into MatLab (The MathWorks, Natick, MA, USA) and run through a custom semi-automated algorithm. In a first step, the algorithm created an average image of the image series. On this average image individual neurons were manually depicted as ROIs. For every ROI at any given time point an average value of fluorescence intensity was determined by averaging the fluorescence values of each pixel within the ROI in every single image of the series, resulting in a calcium trace for every ROI. A baseline region was selected in every trace and measured fluorescence levels of each ROI were converted into relative changes in fluorescence (dF / F_0_) in relation to the selected baseline region. In the next step, the data was transferred to Igor Pro and calcium analysis was conducted with a custom-made program suite. Transients were detected by a peak detection algorithm *[14]*. This algorithm auto-detected calcium transients wherever the amplitude of the calcium signal exceeded 3 - 3.5 SD above the mean, the first derivative crossed zero and the second derivative was negative. 100 % specificity was achieved by increasing this threshold and by calibrating the algorithm using simultaneous juxtacellular recordings. The program suite also enclosed a tool modeling the decay of the calcium deflection and categorizing for exponential and non-exponential decays, therefore taking into account a typical feature of the shape of an AP-related calcium transient. In this tool the decay was fitted by Igor Pro’s built-in curve fitting feature CurveFit, which fits a single exponential curve between the peak and the tail of an autodetected transient. Specificity and sensitivity were confirmed using parallel *in vivo* 2-P GCaMP6f imaging and juxtacellular recordings as described above. The traces were pre-treated by binomial (Gaussian) smoothing 30 - 40 times, followed by a high-pass filter with the end of the reject band at 0.1 / Fs, the start of the pass band at 0.12 / Fs, the number of FIR filter coefficients at 450, and where Fs is the sampling frequency (Hz). The baseline was estimated as a median of 10 s of inactivity one second before the peak. In a further step, the algorithm allowed manual inclusion or exclusion of identified transients. Mean transient frequencies (transients / min) were calculated using the arithmetic mean. Average response latencies upon 2-P optogenetic stimuli as well as average amplitudes of stimulation induced artifacts were determined by calculating arithmetic means. The fraction of responding cells was calculated by dividing the number of responding cells by the number of all stimulated cells. The rate of evoked calcium transients was calculated by dividing the number of evoked calcium transients by the total number of stimulation trials.

### Statistics

Inferential statistics was done with data sets with sample size of n ≥ 5. Statistical significance was tested in GraphPad Prism. Significance levels were * p < 0.05, ** p < 0.01, *** p < 0.001 and **** p < 0.0001. For all data, we first tested for normality using the one-sample Kolmogorov-Smirnov test. In case that the null hypothesis (H_0_) of a normal distribution could not be rejected (for p > 0.05), we employed a parametric test (t test), if H_0_ could be rejected (for p < 0.05) we used a non-parametric test (Mann-Whitney U test). Data sets of Fig. 2 C (number of active cells) and D (number spontaneous calcium transients) where normalized to values at 920 nm (100 %). For significance testing of artifacts in Figure 7 C and E, we tested the mean artifact amplitude calculated as the arithmetic mean of all mean artifacts from calcium traces of every individual cell against the standard deviation of the baseline level. Descriptive statistics given in the text are mean ± s.e.m. and error bars are displayed mean ± s.e.m.. Box-whisker plots are displayed 25 - 75 % (box) and 10 - 90 % percentile (whiskers). All data was quantified in GraphPad Prism and formatted with Adobe Illustrator (Adobe, San Jose, CA, USA).

## Supplementary Materials

Fig. S1. Determining the spatial specificity of 2-P optogenetic stimulations.

Fig. S2. A photostimulation artifact was found in only one highly GCaMP6f-expressing neuron at 1100 nm.

Fig. S3. Using *in vivo* 2-P calcium imaging to identify the current brain state.

## Acknowledgments

We thank Nicolas Ruffini for support in analyzing the photostimulation artifact and Nico Bürger for viral injections.

## Author contributions

T. F., I. A. and J. D. are equally contributing first authors. A. S. designed the study. T. F. performed calcium imaging, all-optical experiments and virus titration. I. A. and H. W. performed cell-attached recordings combined with calcium imaging and optogenetic stimulation. J. D. performed LFP-experiments combined with optic fiber stimulations. T. F., I. A., J. D. and H. W. performed the imaging analysis. I. A. and J. D. performed the analysis of *in vivo* electrophysiology. J. D. performed confocal analysis. I. S. performed PSF measurements. A. S., J. D., T. F., I. A., H. W. and I. S. wrote the manuscript.

## Data and materials availability

All data associated with study will be made available upon request.

## References and Notes

1. Emiliani, V., et al., All-Optical Interrogation of Neural Circuits. J Neurosci, 2015. 35(41): p. 13917–26.

2. Rickgauer, J.P., K. Deisseroth, and D.W. Tank, Simultaneous cellular-resolution optical perturbation and imaging of place cell firing fields. Nat Neurosci, 2014. 17(12): p. 1816–24.

3. Packer, A.M., et al., Simultaneous all-optical manipulation and recording of neural circuit activity with cellular resolution in vivo. Nat Methods, 2015. 12(2): p. 140–6.

4. Carrillo-Reid, L., et al., Imprinting and recalling cortical ensembles. Science, 2016. 353(6300): p. 691–4.

5. Carrillo-Reid, L., et al., Controlling Visually Guided Behavior by Holographic Recalling of Cortical Ensembles. Cell, 2019. 178(2): p. 447–457 e5.

6. Forli, A., et al., Two-Photon Bidirectional Control and Imaging of Neuronal Excitability with High Spatial Resolution In Vivo. Cell Rep, 2018. 22(11): p. 3087–3098.

7. Mardinly, A.R., et al., Precise multimodal optical control of neural ensemble activity. Nat Neurosci, 2018. 21(6): p. 881–893.

8. Zhang, Z., et al., Closed-loop all-optical interrogation of neural circuits in vivo. Nat Methods, 2018. 15(12): p. 1037–1040.

9. Chen, I.W., et al., In Vivo Submillisecond Two-Photon Optogenetics with Temporally Focused Patterned Light. J Neurosci, 2019. 39(18): p. 3484–3497.

10. Marshel, J.H., et al., Cortical layer-specific critical dynamics triggering perception. Science, 2019. 365(6453).

11. Packer, A.M., et al., Two-photon optogenetics of dendritic spines and neural circuits. Nat Methods, 2012. 9(12): p. 1202–5.

12. Yang, W., et al., Simultaneous two-photon imaging and two-photon optogenetics of cortical circuits in three dimensions. Elife, 2018. 7.

13. Busche, M.A., et al., Critical role of soluble amyloid-beta for early hippocampal hyperactivity in a mouse model of Alzheimer’s disease. Proc Natl Acad Sci U S A, 2012. 109(22): p. 8740–5.

14. Arnoux, I., et al., Metformin reverses early cortical network dysfunction and behavior changes in Huntington’s disease. Elife, 2018. 7.

15. Ellwardt, E., et al., Maladaptive cortical hyperactivity upon recovery from experimental autoimmune encephalomyelitis. Nat Neurosci, 2018. 21(10): p. 1392–1403.

16. Buzsaki, G. and K. Mizuseki, The log-dynamic brain: how skewed distributions affect network operations. Nat Rev Neurosci, 2014. 15(4): p. 264–78.

17. Miller, J.E., et al., Visual stimuli recruit intrinsically generated cortical ensembles. Proc Natl Acad Sci U S A, 2014. 111(38): p. E4053–61.

18. Iaccarino, H.F., et al., Gamma frequency entrainment attenuates amyloid load and modifies microglia. Nature, 2016. 540(7632): p. 230–235.

19. Martorell, A.J., et al., Multi-sensory Gamma Stimulation Ameliorates Alzheimer’s-Associated Pathology and Improves Cognition. Cell, 2019. 177(2): p. 256–271 e22.

20. Duda, G.N., et al., Changing the mindset in life sciences toward translation: a consensus. Sci Transl Med, 2014. 6(264): p. 264cm12.

21. Dirnagl, U., Thomas Willis Lecture: Is Translational Stroke Research Broken, and if So, How Can We Fix It? Stroke, 2016. 47(8): p. 2148–53.

22. Dirnagl, U., Rethinking research reproducibility. EMBO J, 2019. 38(2).

23. Piper, S.K., et al., Exact replication: Foundation of science or game of chance? PLoS Biol, 2019. 17(4): p. e3000188.

24. Stosiek, C., et al., In vivo two-photon calcium imaging of neuronal networks. Proc Natl Acad Sci U S A, 2003. 100(12): p. 7319–24.

25. Grienberger, C. and A. Konnerth, Imaging calcium in neurons. Neuron, 2012. 73(5): p. 862–85.

26. Prakash, R., et al., Two-photon optogenetic toolbox for fast inhibition, excitation and bistable modulation. Nat Methods, 2012. 9(12): p. 1171–9.

27. Rochefort, N.L., et al., Sparsification of neuronal activity in the visual cortex at eye-opening. Proc Natl Acad Sci U S A, 2009. 106(35): p. 15049–54.

28. Schmid, F., et al., Assessing sensory versus optogenetic network activation by combining (o)fMRI with optical Ca2+ recordings. J Cereb Blood Flow Metab, 2016. 36(11): p. 1885–1900.

29. Yang, J.W., et al., Optogenetic Modulation of a Minor Fraction of Parvalbumin-Positive Interneurons Specifically Affects Spatiotemporal Dynamics of Spontaneous and Sensory-Evoked Activity in Mouse Somatosensory Cortex in Vivo. Cereb Cortex, 2017. 27(12): p. 5784–5803.

30. Fois, C., P.H. Prouvot, and A. Stroh, A roadmap to applying optogenetics in neuroscience. Methods Mol Biol, 2014. 1148: p. 129–47.

31. Stroh, A., et al., Making waves: initiation and propagation of corticothalamic Ca2+ waves in vivo. Neuron, 2013. 77(6): p. 1136–50.

32. Schwalm, M., et al., Cortex-wide BOLD fMRI activity reflects locally-recorded slow oscillation-associated calcium waves. Elife, 2017. 6.

33. Chen, T.W., et al., Ultrasensitive fluorescent proteins for imaging neuronal activity. Nature, 2013. 499(7458): p. 295–300.

34. Greenberg, D.S., D.J. Wallace, and J.N. Kerr, Imaging neuronal population activity in awake and anesthetized rodents. Cold Spring Harb Protoc, 2014. 2014(9): p. 912–22.

35. Ronzitti, E., et al., Sub-millisecond optogenetic control of neuronal firing with two-photon holographic photoactivation of Chronos. bioRvix, 2016.

36. Iaccarino, H.F., et al., Author Correction: Gamma frequency entrainment attenuates amyloid load and modifies microglia. Nature, 2018. 562(7725): p. E1.

37. Cardin, J.A., et al., Driving fast-spiking cells induces gamma rhythm and controls sensory responses. Nature, 2009. 459(7247): p. 663–7.

