## Supplementary materials for "Navigating the translational roadblock: Towards highly specific and effective all-optical interrogations of neural circuits"

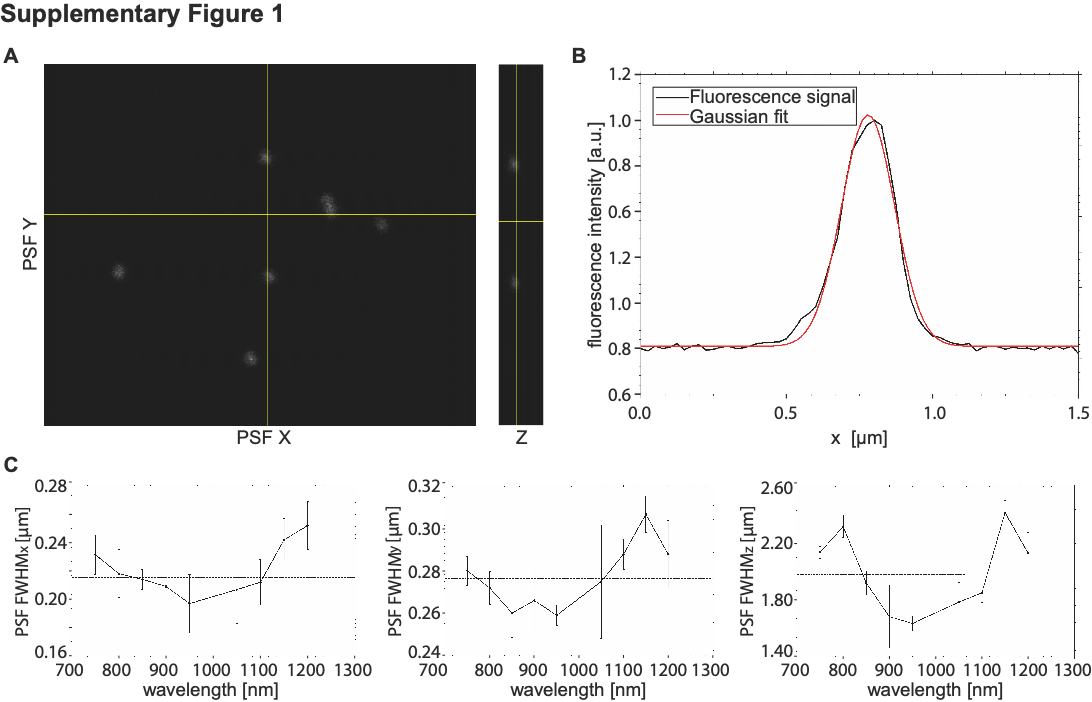

Fig. S1. Determining the spatial specificity of 2-P optogenetic stimulation. A Fluorescent nanobeads, 2-P horizontal (XY) and vertical (YZ) cross sections at 920 nm. B X axis single bead fluorescent signal and gaussian fit for point spread function (PSF) and full width half maximum (FWHM) measurements. C PSF measurements in X, Y and Z direction for varying excitation wavelengths. Error margins are extracted from n = 5 average beads’ measurements for each tested wavelength.

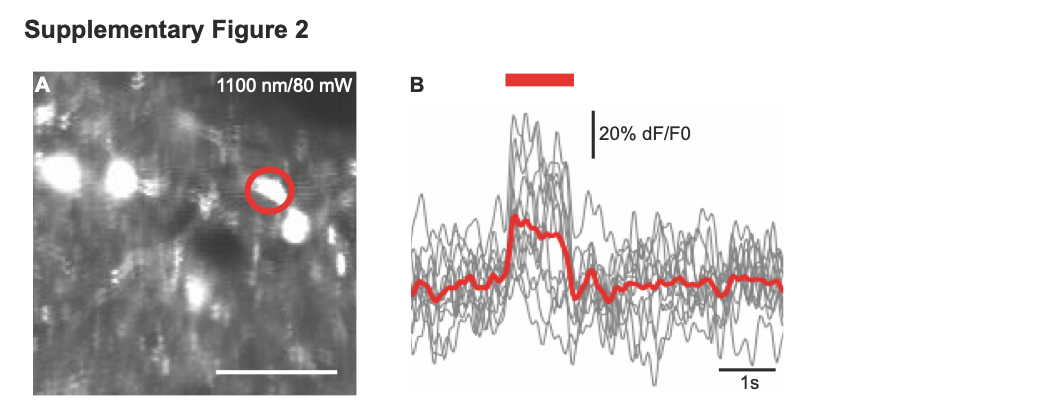
Fig. S2. A photostimulation artifact was found in one highly GCaMP6f-expressing neuron at 1100 nm. A *In vivo* 2-P imaging in layer II/III of mouse visual cortex. Neuron with artifactual signal is depicted with a red circle. Scale bar 50 µm. B Average artifact (red, n = 10 trials) and single trials (grey) upon photostimulation with 80 mW at 1100 nm.

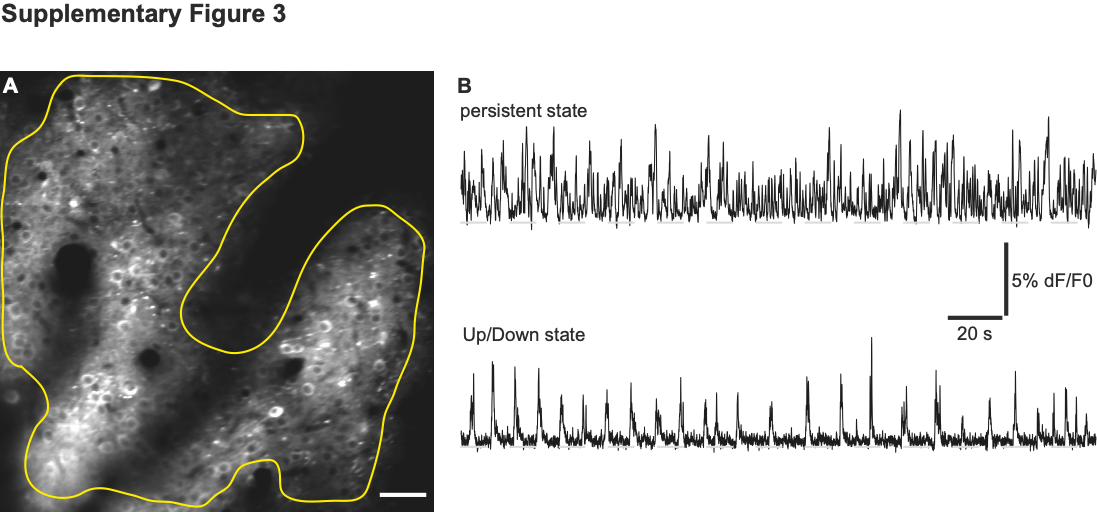
**Fig. S3. Using *in vivo* 2-P calcium imaging to identify the current brain state.** A Typical field of view during 2-P calcium recording of the same cortical microcircuit in layer II/III of mouse visual cortex during different brain states (left). Scale bar 50 µm. B Corresponding calcium traces devised from the entire area of GCaMP6f-expression integrating cellular and neuropil signals. Different brain states are defined by their distinct patterns of spontaneous neural activity and can be induced by varying levels of anesthesia. Light isoflurane anesthesia leads to an activated or persistent state where spontaneous activity is defined by fast low-amplitude oscillations (upper), while deep anesthesia leads to slow oscillations characterized by spontaneous fluctuations between intermittent depolarized Up states and hyperpolarized Down states (lower).
